# Postnatal development of centrifugal inputs to the olfactory bulb

**DOI:** 10.1101/2021.11.15.468595

**Authors:** Johanna K. Kostka, Sebastian H. Bitzenhofer

## Abstract

Processing in primary sensory areas is influenced by centrifugal inputs from higher brain areas, providing information about behavioral state, attention, or context. Activity in the olfactory bulb, the first central processing stage of olfactory information, is dynamically modulated by direct projections from a variety of areas in adult mice. Despite the early onset of olfactory sensation compared to other senses, the development of centrifugal inputs to the olfactory bulb remains largely unknown. Using retrograde tracing across development, we show that centrifugal projections to the olfactory bulb are established during the postnatal period in an area-specific manner. While feedback projections from the piriform cortex are already present shortly after birth, they strongly increase in number during postnatal development with an anterior-posterior gradient. Contralateral projections from the anterior olfactory nucleus are present at birth but only appeared postnatally for the nucleus of the lateral olfactory tract. Numbers of olfactory bulb projecting neurons from the lateral entorhinal cortex, ventral hippocampus, and cortical amygdala show a sudden increase at the beginning of the second postnatal week and a delayed development compared to more anterior areas. These anatomical data suggest that limited top-down influence on odor processing in the olfactory bulb may be present at birth, but strongly increases during postnatal development and is only fully established later in life.

## 1 Introduction

Sensory inputs are a strong driver of neuronal activity in early sensory areas, but sensory processing is not a strict feedforward process. Centrifugal inputs from downstream areas provide information about contextual factors, such as behavioral state, attention, or prior knowledge, that strongly modulate early sensory activity (Gilbert and Sigman, 2007). However, little is known about the development of centrifugal projections.

Development of centrifugal projections to the olfactory bulb (OB) is of particular interest because newborn rodents rely on olfaction for their survival when most other senses are still nonfunctional (Sullivan, 2003; Logan et al., 2012). Odor processing begins with the binding of odor molecules to olfactory receptors on olfactory receptor neurons (ORNs) in the olfactory epithelium. ORNs send direct projections to structures called glomeruli in the OB, the first central processing stage for olfactory information. In the glomeruli, ORNs synapse onto dendrites of mitral and tufted cells, the principal cells in the OB, that transmit olfactory information to a range of brain areas, including the anterior olfactory nucleus (AON), piriform cortex (PIR), olfactory tubercle, nucleus of the lateral olfactory tract (nLOT), cortical amygdala (CoA), and lateral entorhinal cortex (LEC) (Igarashi et al., 2012). In the adult brain, most of these areas, with exception of the olfactory tubercle, send glutamatergic feedback projections to the OB, providing fast top-down modulation of olfactory processing (Luskin and Price, 1983; Shipley and Adamek, 1984; Padmanabhan et al., 2019; Zandt et al., 2019). Additionally, glutamatergic feedback from CA1 of the ventral hippocampus to the OB has been described in adult mice (Padmanabhan et al., 2019).

In adults, centrifugal projections to the OB mainly target inhibitory neurons in the glomerular layer and granule cell layer and are thereby ideally positioned to modulate network activity (Boyd et al., 2012; Markopoulos et al., 2012). Centrifugal inputs to OB provide diverse feedback critical for the formation of odor-reward associations (Kiselycznyk et al., 2006; Gao and Strowbridge, 2009; Markopoulos et al., 2012; Boyd et al., 2015). The ability of rodents to form odor-reward associations early in life suggests that feedback projections to the OB may be established early during development (Logan et al., 2012). While feedforward projections from the OB are established at birth (Walz et al., 2006) and OB activity drives downstream areas early in life (Gretenkord et al., 2019; Kostka et al., 2020; Kostka and Hanganu-Opatz, 2021), the development of centrifugal projections to the OB is largely unknown. We took advantage of retrograde virus-labeling to investigate the maturation of glutamatergic centrifugal inputs to the main OB during postnatal development in mice.

## 2 Materials and Methods

### 2.1 Animals

All experiments were performed in compliance with the German laws and the guidelines of the European Union for the use of animals in research (European Union Directive 2010/63/EU) and were approved by the local ethical committee (Behörde für Gesundheit und Verbraucherschutz Hamburg, ID 15/17).

Experiments were carried out in C57BL/6J mice of both sexes. Timed-pregnant mice from the animal facility of the University Medical Center Hamburg-Eppendorf were housed individually at a 12□h light/12□h dark cycle and were given access to water and food ad libitum. The day of birth was considered postnatal day (P) 0.

### 2.2 Virus injections

For retrograde labeling of OB-projecting neurons, C57BL/6J mice received unilateral injections of AAVrg-CaMKIIα-mCherry (200 nl at 200 nl/min, titer 2×10^13^ vg/ml, #114469-AAVrg, Addgene, MA, USA) into the right main OB (0.5 mm lateral from midline, 0.5 mm rostral to the inferior cerebral vein, 0.5-1.0 mm deep). Injections were performed at P0, P3, P6, P9, P12, or P49 in a stereotaxic apparatus using a micropump (Micro4, WPI, Sarasota, FL) under anesthesia. Following injection, the syringe was left in place for >60 s to reduce reflux. Mice were kept on a heating blanket until full recovery from anesthesia and returned to their home cage.

9 days after virus injection, mice were transcardially perfused with 4% paraformaldehyde (PFA). Brains were removed, post fixed in PFA for 24-48 hours, and stored in phosphate buffered saline (PBS) with 0.02 sodium azide. Brains were sliced into coronal sections at 100 μm and mounted with Vectashield with DAPI (Vector Laboratories). Fluorescence images were taken to validate injections sites and to identify areas with retrogradely labeled neurons.

### 2.3 Cell quantification

Single images (2048 × 2048 pixels) were taken with a confocal microscope (Zeiss, Germany) using a 20x objective with a 405 nm laser for DAPI and a 568 nm laser for mCherry. This resulted in pixel size of 0.16 μm^2^, corresponding to images of 319.5 μm^2^. Brain areas were identified according to the Mouse reference atlas from Allen brain atlas (Lein et al., 2007). Cells were detected with Cellpose (Stringer et al., 2021), a deep learning-based cellular segmentation algorithm in Python 3.8. Parameters were kept constant for all images and results were validated by visual inspection. Data were imported and analyzed in Matalab R2021a. Data are shown as mean ± standard error of the mean (SEM).

## 3 Results

### 3.1 Retrograde labeling of OB-projecting neurons across development

To investigate the development of centrifugal projections to the OB, we injected AAVrg-CaMKIIα-mCherry in the right OB of P0 (n=3), P3 (n=2), P6 (n=2), P9 (n=6), P12 (n=5), or P49 (n=4) mice to transduce neurons with axons projecting to the injection area at the day of injection (Figure 1A). P49 mice were considered adult since we did not assume further changes in OB-feedback projections at that age. Mice were perfused 9 days after injection to allow for the expression of the plasmid. Injection areas were confirmed post mortem (Figure 1B, C). Although injection areas in the main OB were carefully inspected, due to the retrograde labeling of neurons in nearby areas we cannot rule out completely that some injections may have extended into neighboring areas. Few brains with labeled neurons in contralateral OB or orbitofrontal cortex, which indicates that injections were not limited to the OB, were excluded from the analysis. As a side note, we observed mCherry expression in OB mitral cells and granule cells during development indicating expression of CaMKIIα in both cell types, similar to other studies (Liu, 2000; Shani-Narkiss et al., 2020), but in contrast to a study reporting that only granule cells in OB would express CaMKIIα (Zou et al., 2002).

**Figure 1.**
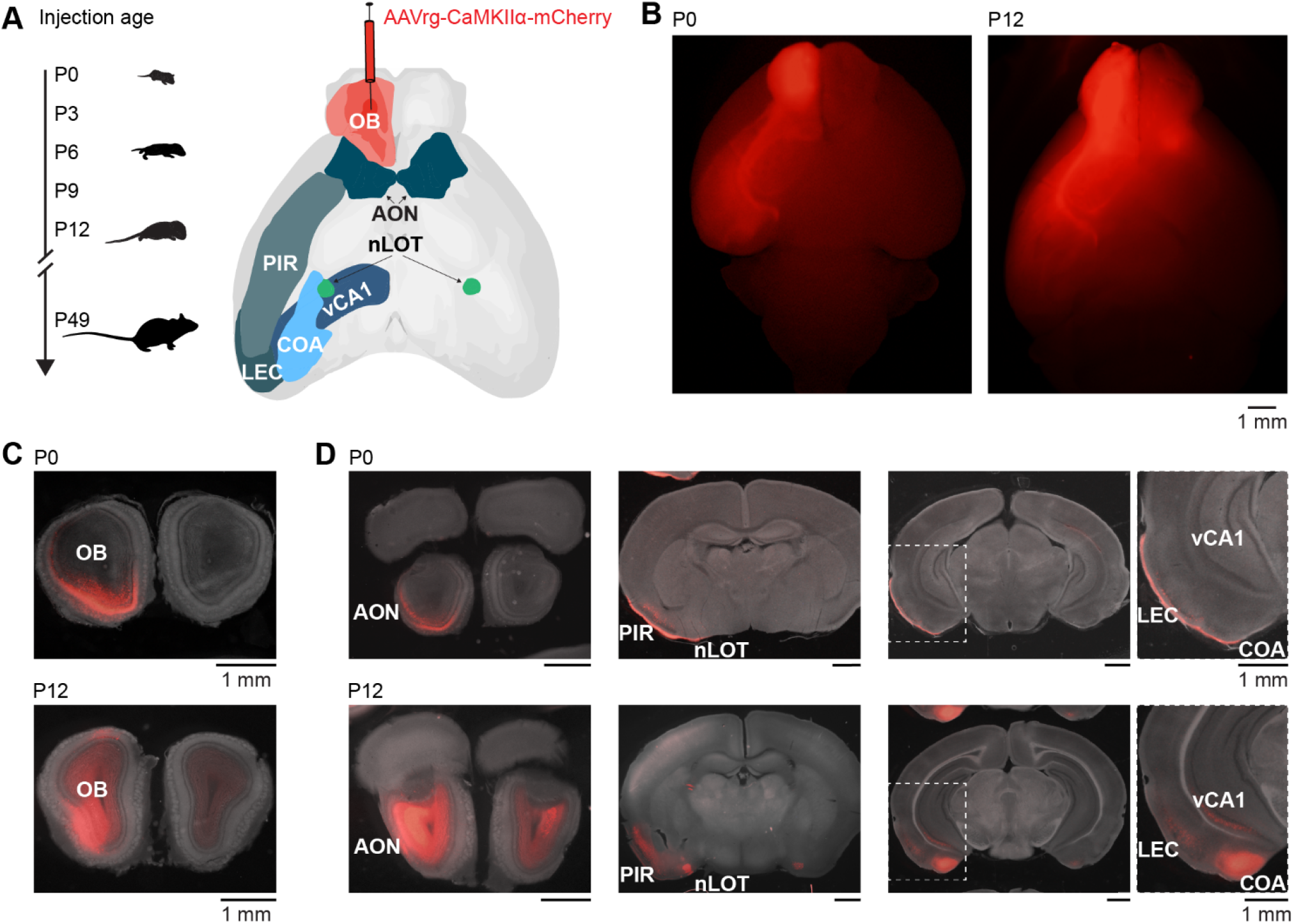
Retrograde tracing to investigate the development of centrifugal projections to OB. **(A)** Scheme illustrating the investigation of glutamatergic centrifugal projections to the OB across development. Mice were injected with the retrograde virus AAVrg-CaMKIIα-mCherry into the right OB at P0, P3, P6, P9, P12, or P49 and perfused 9 days after injection **(B)** Representative whole-brain fluorescence images from ventral view of mCherry expression in LOT and retrogradely labels neurons for unilateral OB injections at P0 and P12. **(C)** Representative fluorescence images of mCherry expression in coronal OB slices at the injection site for OB injections at P0 and P12. **(D)** Representative fluorescence images of retrogradely labeled mCherry expression in AON, nLOT, PIR, CoA, LEC, and vCA1 for mice shown in C.

Brain slices were visually inspected in a fluorescence microscope for mCherry expression. Expression was found in an age-dependent manner bilaterally in AON and nLOT, and ipsilaterally in PIR, CoA, LEC, and CA1 of the ventral hippocampus (vCA1) (Figure 1D). As previously reported, no OB-projecting neurons were found in the olfactory tubercle (Zandt et al., 2019). Of note, CaMKIIα is mainly expressed in glutamatergic neurons, thus neuromodulatory inputs to the OB were not considered in this study, but have been described in adults (Brunert and Rothermel, 2021).

### 3.2 Development of OB-projecting neurons in bilateral AON and nLOT

We took confocal images from retrogradely labeled areas and mCherry expressing cells were counted automatically in images with a size of 319.5 μm^2^ with Cellpose (Stringer et al., 2021) followed by visual confirmation (Figure 2A). Automatic detection worked equally well for the different ages. Similar to the adult brain, bilateral projections from AON and nLOT to the OB were found during development (Figure 2B). OB-projecting neurons in AON were present already at birth at low numbers and gradually increased in number with age (Figure 2C). At P12, numbers of OB-projecting neurons had already reached 72% and 58% of adult (P49) levels for ipsilateral and contralateral AON, respectively (Figure 2D). Similar numbers of OB-projecting neurons were found in ipsilateral and contralateral AON from P0 to P9 but were higher for ipsilateral AON at older age.

**Figure 2.**
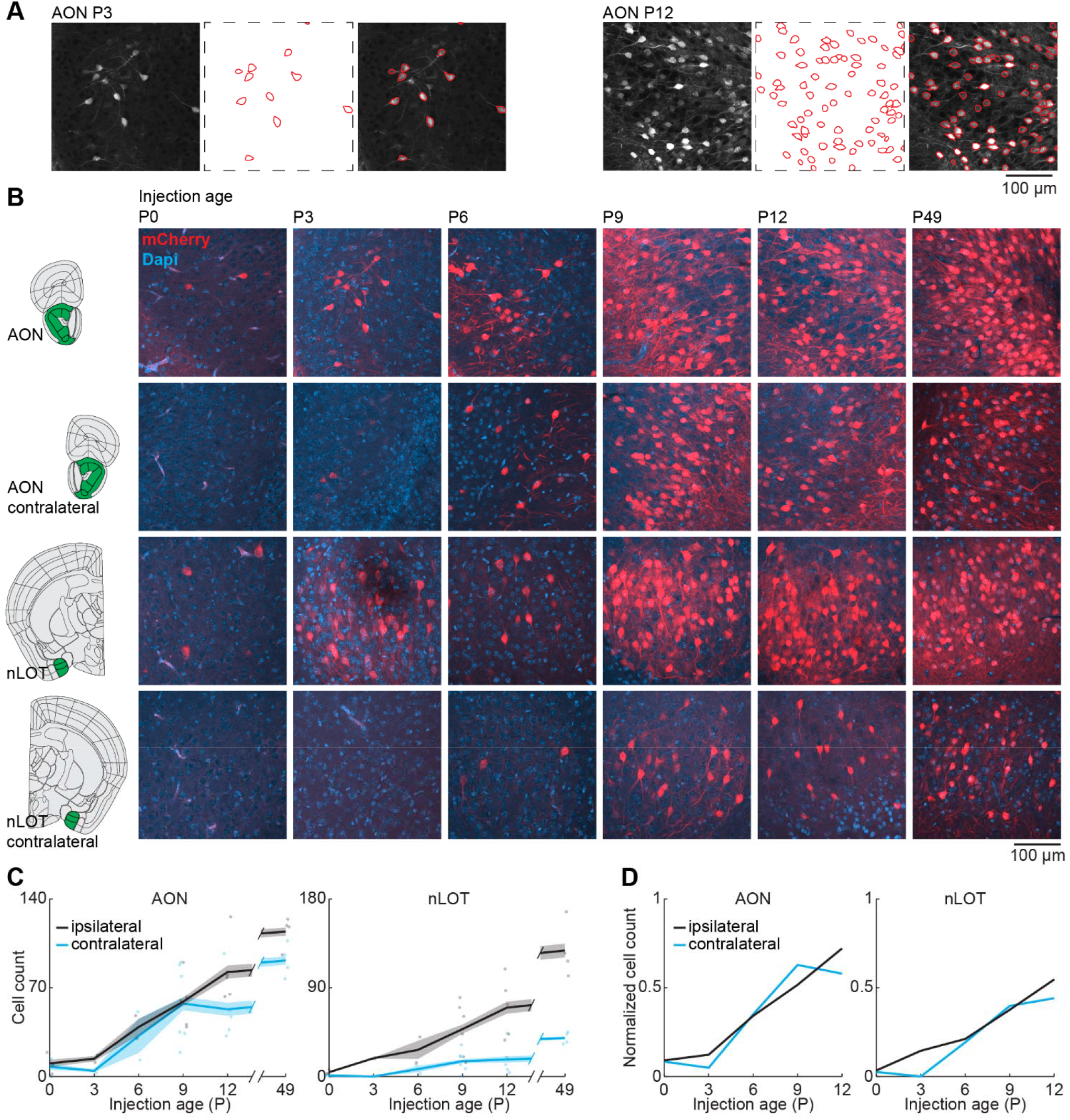
Bilateral centrifugal projections to OB from AON and nLOT. **(A)** Representative confocal images, detected cell outlines (red), and their overlay image for AON after OB injection at P3 and P12. **(B)** Representative confocal images of retrogradely labeled cells in ipsilateral and contralateral AON and nLOT after injection of retrograde virus into the right OB at P0, P3, P6, P9, P12, or P49. Reference images are from the Allen brain reference atlas for adult mice (Lein et al., 2007). **(C)** Quantification of retrogradely labeled cells in ipsilateral and contralateral AON and nLOT across development. Cell numbers were counted in confocal images of 319.5 μm2. **(D)** Average number of retrogradely labeled cells in ipsilateral and contralateral AON and nLOT across development normalized to adult levels at P49.

AON is very close to OB, so we investigated if centrifugal projections from the more posterior nLOT are also present at birth. Similar to AON, centrifugal projections from ipsilateral nLOT were present at birth and increased gradually with age (Figure 2C). However, the number of centrifugal projection neurons in contralateral nLOT was lower and developed later, starting to be reliably detected at P6. Compared to adult mice, the numbers of OB-projecting neurons at P12 in ipsilateral and contralateral nLOT were at 55% and 44%, respectively.

### 3.3 Gradual development of OB feedback projections from PIR

Next, we looked at the development of centrifugal projections from the ipsilateral PIR to the OB. The PIR stretches over a substantial part of the brain in anterior-posterior position. To cover the full extent of PIR, we quantified the number of OB-projecting neurons in the anterior (aPIR), intermediate (iPIR), and posterior (pPIR) part of the PIR (Figure 3A). Across development, we saw an anterior-posterior gradient in the number of labeled neurons with most OB-projecting neurons in the anterior part of PIR (Figure 3B). This gradient persisted into adulthood (P49). However, already at birth, we found OB-projecting neurons in all three parts of the PIR, and the numbers of labeled neurons gradually increased with age in all three parts. At P12 numbers of OB-projecting neurons were already at 77% of adult levels, but only at 62% for iPIR and 39% for pPIR (Figure 3C). Thus, centrifugal projections to the OB from more anterior parts of the PIR are not only higher in numbers but also develop earlier compared to more posterior parts.

**Figure 3.**
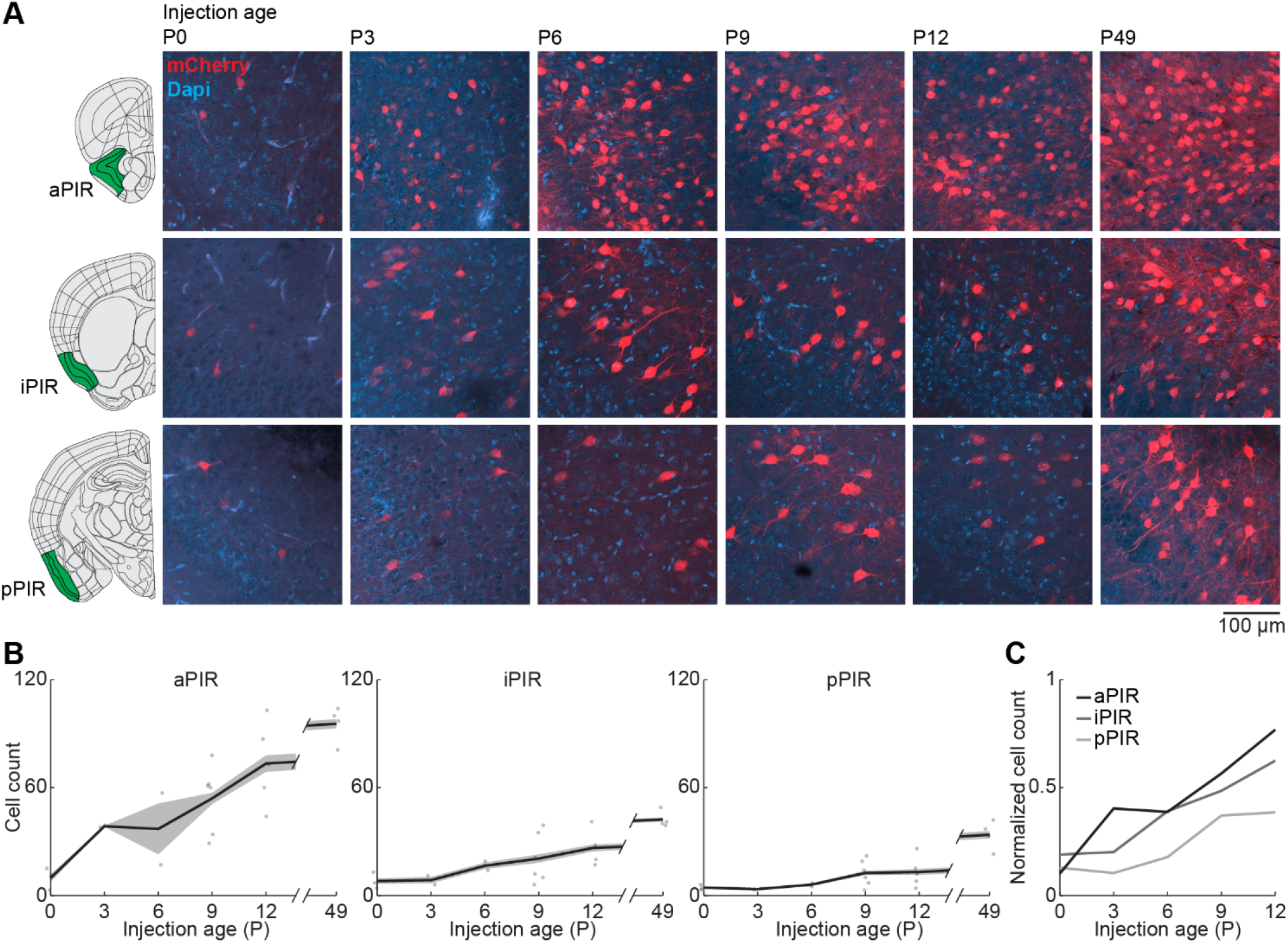
Centrifugal input from PIR develops early and increases with age. **(A)** Representative confocal images of retrogradely labeled cells in anterior, intermediate, and posterior PIR after injection of retrograde virus into the right OB at P0, P3, P6, P9, P12, or P49. Reference images are from the Allen brain reference atlas for adult mice (Lein et al., 2007). **(B)** Quantification of retrogradely labeled cells in anterior, intermediate, and posterior PIR across development. Cell numbers were counted in confocal images of 319.5 μm^2^. **(C)** Average number of retrogradely labeled cells in anterior, intermediate, and posterior PIR across development normalized to adult levels at P49.

### 3.4 Abrupt increase in OB-projecting neurons in posterior brain areas from P6 to P9

Finally, we looked at centrifugal inputs to OB from more posterior areas CoA, LEC, and vCA1, previously described to send ipsilateral projections to the OB in adult mice (Padmanabhan et al., 2019) (Figure 4A). At birth, OB-projecting neurons were absent from CoA and vCA1, and very few neurons were labeled in LEC (Figure 4B). Numbers of labeled neurons stayed absent/low during the first postnatal week but suddenly increased from P6 to P9 for all three areas. The numbers of OB-projecting neurons in CoA were higher than in LEC and vCA1 at all ages investigated. At P12, numbers of OB-projecting neurons were at 47% of adult levels (P49) for CoA and at 37% for LEC, but only at 15% for vCA1, indicating a late development of centrifugal projections from vCA1 to OB (Figure 4C).

**Figure 4.**
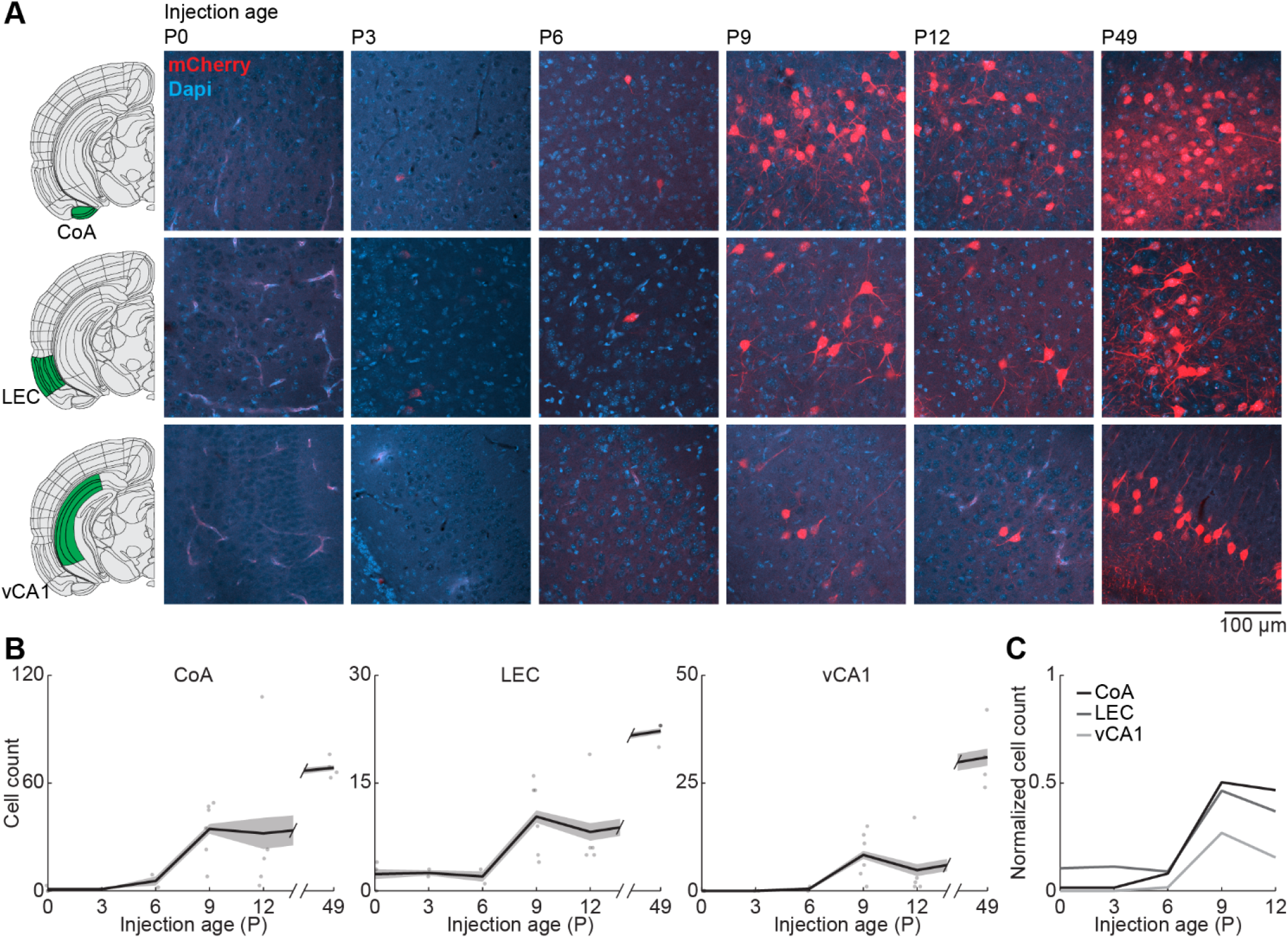
Centrifugal input from posterior brain areas develops in the second postnatal week. **(A)** Representative confocal images of retrogradely labeled cells in CoA, LEC, and vCA1 after injection of retrograde virus into the right OB at P0, P3, P6, P9, P12, or P49. Reference images are from the Allen brain reference atlas for adult mice (Lein et al., 2007). **(B)** Quantification of retrogradely labeled cells in CoA, LEC, and vCA1 across development. Cell numbers were counted in confocal images of 319.5 μm2. **(C)** Average number of retrogradely labeled cells in CoA, LEC, and vCA1 across development normalized to adult levels at P49.

## 4 Discussion

We performed retrograde virus-labeling across development to describe the formation of centrifugal projections to the OB. We found that centrifugal inputs from AON, nLOT, and PIR are already present at birth, but increase extensively in a gradual manner during postnatal development. Contralateral inputs from AON were present at birth, but were not detected before P6 for nLOT. Centrifugal inputs from PIR develop in an anterior-posterior gradient with OB-projecting neurons from anterior parts developing earlier and reaching higher numbers. Centrifugal inputs from CoA and vCA1 only started to be detected at P6 whereas few retrogradely labeled neurons were present at birth in LEC. The development of OB-projecting neurons in areas related to emotional and memory processing such as CoA, vCA1, and LEC was characterized by a sudden increase at the start of the second postnatal week.

Viral injections of the same volume and concentration were used for all age groups, despite the change in brain size, and may have resulted in a smaller relative injection area in the OB for older age groups. We expected to see an increase of retrogradely labeled neurons with age and therefore decided to keep injection parameters constant to make sure that increases with age are not artificially induced by adapting injection volumes to brain size. Thus, the actual age-related increase of OB-projecting neurons seen for all areas may be slightly underestimated.

In the adult brain, the OB also receives inputs from neuromodulatory areas, such as noradrenergic input from the locus coeruleus, serotonergic input from the raphe nuclei, and cholinergic input from the basal forebrain (Brunert and Rothermel, 2021). These inputs have been implicated in the modulation of odor discrimination and odor learning. In this study we focused on glutamatergic inputs to the OB, but a similar approach can be used in future studies to address the development of neuromodulatory inputs and their relevance for odor-driven behavioral abilities early in life.

Olfactory information is processed in two stages in the OB: at the glomerular level, where local interneurons mediate inhibition within and between glomeruli and through lateral and recurrent inhibition of mitral and tufted cells by inhibitory interneurons such as granule cells in the external plexiform layer (Nagayama et al., 2014). Centrifugal inputs to the OB mainly target inhibitory neurons in the glomerular and granule cell layer of the OB but have only weak direct inputs onto mitral cells (Boyd et al., 2012; Markopoulos et al., 2012). Thereby, centrifugal inputs to OB are ideally positioned to modulate olfactory processing and coordinated network activity in the olfactory system. Interestingly, the generation and maturation of inhibitory neurons in OB extends well into the postnatal period (Batista-Brito et al., 2008).

In adult rodents, coherent beta oscillations between OB and brain areas such as PIR, LEC, hippocampus, and prefrontal cortex have been implicated in memory processing and decision making (Martin et al., 2006, 2007; Igarashi et al., 2014; Symanski et al., 2021). Interestingly, the generation of beta oscillations in OB has been shown to depend on centrifugal inputs (Neville and Haberly, 2003; Ravel et al., 2003). Previous studies have shown that neuronal activity in OB drives oscillatory activity in the beta frequency range in LEC, hippocampus, and prefrontal cortex already at the beginning of the second postnatal week (Kostka and Hanganu-Opatz, 2021). Considering the emergence of feedback projections to the OB from memory-related brain areas around the same time period suggests that already at this age past experience could shape sensory processing. The area-specific development of centrifugal projections to the OB suggests that top-down modulation changes with age. We show that feedback from early olfactory areas, such as AON and aPIR, develops first and presumably contributes to basic sensory processing already shortly after birth. In contrast, the delayed maturation of feedback from higher brain areas, such as CoA, LEC and vHP, suggests that valence and memory dependent modulation of OB activity only emerges later in life. Further research is required to understand the functional role of area-specific centrifugal inputs for olfactory processing during neonatal development.

## 6 Author Contributions

J.K.K. and S.H.B. designed the study and performed the experiments. J.K.K. and S.H.B. analyzed the data. J.K.K. and S.H.B. interpreted the data and wrote the manuscript.

## 8 Acknowledgments

We thank Dr. Ileana L. Hanganu-Opatz for conceptual and financial support. We thank A. M. Thies, A. Marquardt, P. Putthoff, and A. Dahlmann for excellent technical assistance.

## 9 Data Availability Statement

The datasets generated for this study are available from the corresponding author on request.

